# Quantifying replication stress in cancer without proliferation confounding

**DOI:** 10.1101/2025.04.10.648133

**Authors:** Philipp Jungk, Maik Kschischo

## Abstract

Replication stress (RS) is a major driver of genomic instability and cancer development, caused by impaired DNA replication that can lead to chromosomal instability (CIN). Although RS is mechanistically linked to CIN, its relationship with cellular proliferation is complex. Depending on the context, RS can either promote or suppress cell growth. Existing RS gene expression signatures overlook this complexity, relying on the overexpression of oncogenes such as MYC, which introduces a proliferation bias.

To disentangle genuine RS from confounding cell cycle and proliferation transcription profiles, we developed and validated a novel gene expression signature that accurately predicts RS independently of oncogene activity. This signature captures RS-related transcriptional changes across diverse cellular contexts, enabling a more robust and proliferation-independent measure of RS in both experimental and clinical samples.

Applying our signature to patient data, we discovered a link between RS and the non-homologous end-joining (NHEJ) DNA repair pathway. Specifically, we observed that MSH2 and MSH6—core components of mismatch repair—are associated with elevated RS and may indicate a shift toward NHEJ-mediated repair under stress conditions.

Our study provides a refined approach to quantify RS and sheds light on its broader impact on DNA repair network dynamics.

**Author summary:** Cancer cells often display defects in faithful DNA replication, which can result in permanent chromosomal alterations during cell division. One major cause of such damage is underreplicated DNA resulting from replication stress (RS). RS describes the slowing or stalling of DNA replication due to factors like a premature start of replication, before necessary replication factors are assembled. This makes RS a key driver in cancer development. Current methods quantifying RS rely heavily on overexpression of oncogenes that deregulate the cell cycle leading to an increase in cell growth and confounding the identification of RS.

To address this, we carefully selected samples displaying RS from diverse sources and identified genes biologically and statistically linked to RS, independent of cell growth signals. Using this new signature, we found a link between RS and a low-fidelity DNA repair mechanism. By improving RS quantification, our work could provide new insights into how cells respond to replication stress and therefore lead to improved methods for diagnosing and treating cancers linked to RS.

## Introduction

Replication stress (RS) is a significant contributor to genomic instability and cancer development. It is defined as slowed or stalled DNA replication caused by factors like unusual DNA structures, impaired origin licensing, nucleotide pool depletion and transcription-replication collisions [1]. These disruptions stall or collapse replication forks, leading to under-replicated DNA, double-strand breaks (DSB) and ultimately to structural chromosomal instability (S-CIN). RS also links S-CIN and whole chromosomal instability (W-CIN), describing the continuous gain or loss of entire chromosomes [2], through mitotic errors caused by mechanisms like impaired kinetochore attachment and abnormal spindle assembly [3–5]. Chromosomal instability (CIN), a hallmark of cancers, is thought to promote tumor growth [6] by deregulating pathways via gains in oncogenes (e.g. *CCNE1*, *MYC*) or losses in tumor suppressors (e.g. *PTEN*, *RB1*) (7,8).

Accurately quantifying RS based on omics-data is essential for linking it to other tumor-related processes, translating findings from biological experiments into primary tumor databases, and informing prognosis and therapy. In cell line experiments RS is commonly induced by drug treatment (e.g. aphidicolin (APH) or hydroxyurea), oncogene overexpression (e.g. *MYC*, or *CCNE1*), or gene knockouts (e.g. *DHX9*) (9– 11).

While effective in controlled laboratory settings, each method also alters cellular programs unrelated to RS, skewing phenotypic correlations if examined in isolation. For instance, oncogene overexpression may bias the transcriptome toward genes linked to premature cell cycle progression, potentially obscuring RS-specific effects like the activation of DSB repair pathways in S-phase [1,12–14]. Despite these limitations, current gene expression-based RS quantifications emphasize oncogene-related phenotypes [15,16] likely due to their established association to RS in cell lines and frequent natural occurrence in primary tumors [10].

CIN was suggested to paradoxically affect proliferation, enhancing fitness in some context while impairing it in others [6,17]. Further studies have extended upon these discussions, including evidence of a similar paradoxical relationship between RS and proliferation. For instance, CIN signatures were shown to positively correlate with the proliferation markers like *PCNA* and *MKI67*, as well as proliferation scores in primary tumors, suggesting that genomic instability might drive tumor growth in advanced stages of cancers [17–19]. CIN, assessed through comparative genomic hybridization and image cytometry, has been linked to gene expressions indicating cancer development and cellular growth in breast cancers [20]. In mouse models and murine embryonic fibroblasts, 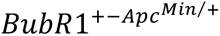 induced CIN led to the development of higher growth rates [21]. Similarly, a causal link to proliferation is suggested by nucleotide depletion in highly proliferative cells inducing RS [22,23]. However, evidence for this positive association in cell lines is limited. Experiments suggesting that genomic instability increases proliferation required additional defects in cell cycle regulators (e.g. *ATR*, *ATM*), alongside oncogene-induced RS [24,25]. Inhibiting the checkpoint assuring faithfully replicated DNA in oncogene-driven proliferation environments may increase proliferation and overshadowing CIN/RS-dependent effects. Further experiments in cell lines do suggest that CIN and RS generally reduce cellular fitness. For example, even low RS levels induced by 100 nM APH were shown to decrease proliferation in cell lines [26], while increased karyotype heterogeneity in NCI60 cells correlates with slower doubling times [17]. Gene set enrichment analysis (GSEA) of CIN-associated genes in breast cancer cell lines indicated reduced cell cycle activity and a shift of their transcriptome toward a mesenchymal state [17,27]. Interestingly, some mouse models harboring CIN were shown to reduce proliferation. For instance, 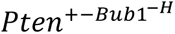 mice exhibited increased abnormal chromosomal numbers, while *PCNA*-positive cells were reduced compared to *Pten*^+−^ mice with stable chromosome numbers [28]. Another study found a simultaneous decrease in expression of the proliferation marker *Ki67* and an increase in levels of DNA damage as well as aneuploidy in *SA1*-deficient mice [29]. While most studies on the association of genomic instability and proliferation focus on CIN, the limited evidence on RS displaying similar trends, suggests that the CIN-proliferation relationship extends to RS [2,30–32].

To determine a suitable quantification of RS, we first systematically evaluate existing RS scores and analyze their association with a chromosomal complexity score while accounting for the influence of other cancer hallmarks. Using public gene expression data from various RS-inducing methods, we develop a new RS signature that performs well even on unseen experimental data, while minimizing the confounding effects of oncogene overexpression and capturing the complex relationship between proliferation to RS. Finally, we apply this novel RS signature to large genomic databases to uncover a proteomic link to the non-homologous end-joining pathway, demonstrating its utility for uncovering new insights into cancer biology.

## Results

### Existing signatures fail to predict replication stress in cancer cell lines

To estimate the performance of previously published replication stress signatures, we gathered public RNA-sequencing data from various studies using different methods for replication stress induction (Table 1): Three of the methods used RS-inducing oncogene overexpression (*CCNE1*, *CDC25A*, *MYC*) [33,34], while three additional methods induced RS by silencing genes affecting RS-pathways (*DHX9*, *CDA*, *SMAD4*) [11,35]. Samples were divided into those with and without replication stress.

**Table 1:** Samples with gene amplification or knockout to induce replication stress.

| *Study_ID* | *Cell line* | *RS Technique* | *GEO series* |
| --- | --- | --- | --- |
| *GSE171845* | RPE_1 | CCNE1 | GSE171845 |
| *GSE185512* | RPE_1 | CDC25A, CCNE1, MYC | GSE185512 |
| *GSE185512* | RPE_1_NOp53 | CDC25A, CCNE1, MYC | GSE185512 |
| *GSE185512* | BT549 | CDC25A, CCNE1, MYC | GSE185512 |
| *GSE185512* | MDA_MB231 | CDC25A, CCNE1, MYC | GSE185512 |
| *GSE185512* | HCC1806 | CDC25A, CCNE1, MYC | GSE185512 |
| *GSE244103* | H446 | DHX9_KO | GSE244103 |
| *GSE244103* | H82 | DHX9_KO | GSE244103 |
| *GSE244103* | H196 | DHX9_KO | GSE244103 |
| *GSE252544* | Mia_PACA_2 | CDA_KO | GSE252544 |
| *GSE292334* | HCT116 | SMAD4_KO | GSE292334 |
| *GSE292334* | HT29 | SMAD4_KO | GSE292334 |

We applied two published RS signatures to estimate the replication stress in the dataset’s samples: (i) The repstress score based on *MYC* overexpression, DNA damage checkpoint activation and sensitivity of checkpoint inhibitors and (ii) the oncoRS score based on oncogene overexpression in cell lines and oncogene amplification in primary tumors [15,16]. Notably, samples with and without RS-induction displayed no difference in the repstress scores, indicating that the repstress score was not able to predict RS status (Figure 1A), while the oncoRS signature was only able to differentiate RS from non-RS in samples with *MYC*-based RS-induction (Figure 1B).

**Fig 1:**
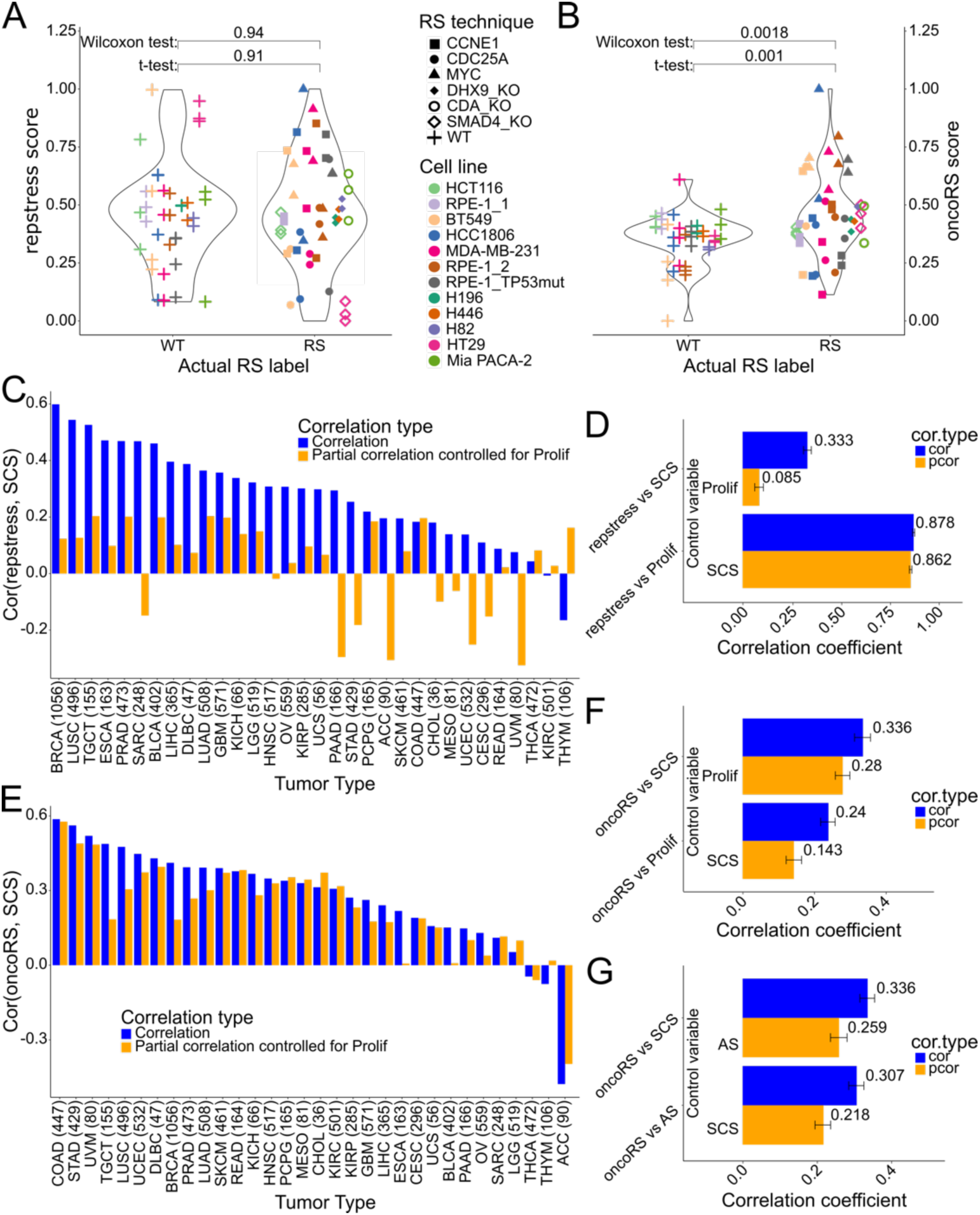
Assessment of RS prediction scores and their robustness across multiple studies and associated phenotype scores. Evaluation of two replication stress scores on a dataset including six different ways of inducing replication stress in multiple cell lines. The y-axis represents the replication stress levels as predicted by the repstress score **(A)** and the oncoRS score **(B)**, while the x-axis distinguishes between the actual status of samples either with replication stress (RS) or without (WT). **(C)** Spearman correlation of the repstress score and the SCS within tumors of TCGA. The y-axis visualizes the non-adjusted (blue) correlation coefficient and the partial (orange) correlation coefficient between repstress score and SCS accounting for proliferation in each tumor type (x-axis). **(D)** Pan-cancer correlation between repstress score and SCS and proliferation with (orange) and without (blue) correction for confounding effects of the annotated control variable using partial correlation. The y-axis shows the comparison pair, while the x-axis represents the (partial) correlation coefficient, which is additionally written next to the bar. **(E)** Spearman correlation of the oncoRS score and the SCS within tumors of TCGA. The y-axis visualizes the non-adjusted (blue) correlation coefficient and the partial (orange) correlation coefficient between oncoRS score and SCS accounting for proliferation in each tumor type (x-axis). **(F),(G)** Pan-cancer correlation between oncoRS score and SCS, proliferation **(F)** and aneuploidy **(G)** with (orange) and without (blue) correction for confounding effects of the annotated control variable using partial correlation. The y-axis shows the comparison pair, while the x-axis represents the (partial) correlation coefficient, which is additionally written next to the bar. **(D),(F),(G)** Error bars represent the 95% confidence interval. SCS: Structural Complexity Score; TCGA: The Cancer Genome Atlas; AS: Aneuploidy Score; Prolif: proliferation

To further assess the physiological reliability of these signatures to predict RS, we used the structural complexity score (SCS), which quantifies the heterogeneity of a sample’s copy number landscape to approximate S-CIN [2]. Further we used a gene expression-based quantification of the proliferation rate [36] and an aneuploidy score based on the absolute number of arm-level alterations of a sample [37].

We evaluated the association between RS scores and SCS in primary tumors of The Cancer Genome Atlas (TCGA) by calculating the Spearman correlation. Using partial correlation, we investigated the confounding of this RS-SCS association by proliferation, potentially resulting from bias toward oncogene and cell cycle activities.

The repstress score displayed a strong correlation with proliferation and a significant association with the SCS within (Figure 1C) and across (Figure 1D) TCGA tumor types. However, this association with the SCS was significantly reduced when controlling for proliferation. Additionally, the repstress signature genes are cell cycle genes with a strong correlation to the proliferation score and gene expressions of the proliferation markers *PCNA* and *MKI67* (Supplementary Table 1). In combination this suggests that the repstress score-SCS association is largely mediated by proliferation.

The oncoRS score’s correlation with the SCS remained largely unaffected after accounting for proliferation (Figure 1E, F). Since the oncoRS score is partially based on oncogene amplification, we further investigated the signature for confounding by arm-level aneuploidies. A pan-cancer comparison of oncoRS and aneuploidy revealed a similar correlation as with the SCS. Partial correlation showed that the correlation of oncoRS and the S-CIN measure may be influenced by aneuploidies, as it resulted in a mild reduction in the oncoRS-SCS association (Figure 1G). Independent validation of the oncoRS signature genes remains limited with *NAT10* being the only gene confirmed to display an association with RS markers in tumors.

### Moderate APH dosage leads to a non-physiological RS response with an accumulation of cells in S-phase

We aimed to derive a replication stress signature without intrinsic biases toward proliferation, CNA, or oncogene-related processes. To achieve this, we expanded the dataset in Table 1 to include drug-induced RS using APH and hydroxyurea (Supplementary Table 2). In our systematic approach (Figure 2A), we first calculated differentially expressed genes in RS samples using a bootstrap approach of the limma-voom pipeline [38,39]. To further reduce the dimensionality of the selection, we integrated prior biological knowledge by including only genes involved in any processes potentially associated with RS, such as DNA replication and repair among others (Supplementary Table 3). The top-ranked genes were then selected for the signature. In the last step, batch corrected log-expressions of the signature genes were used to train a regularized logistic regression model. To assess the model’s performance, the estimated coefficients for the genes were used to calculate RS scores in an unseen test dataset (Figure 2B).

**Fig 2:**
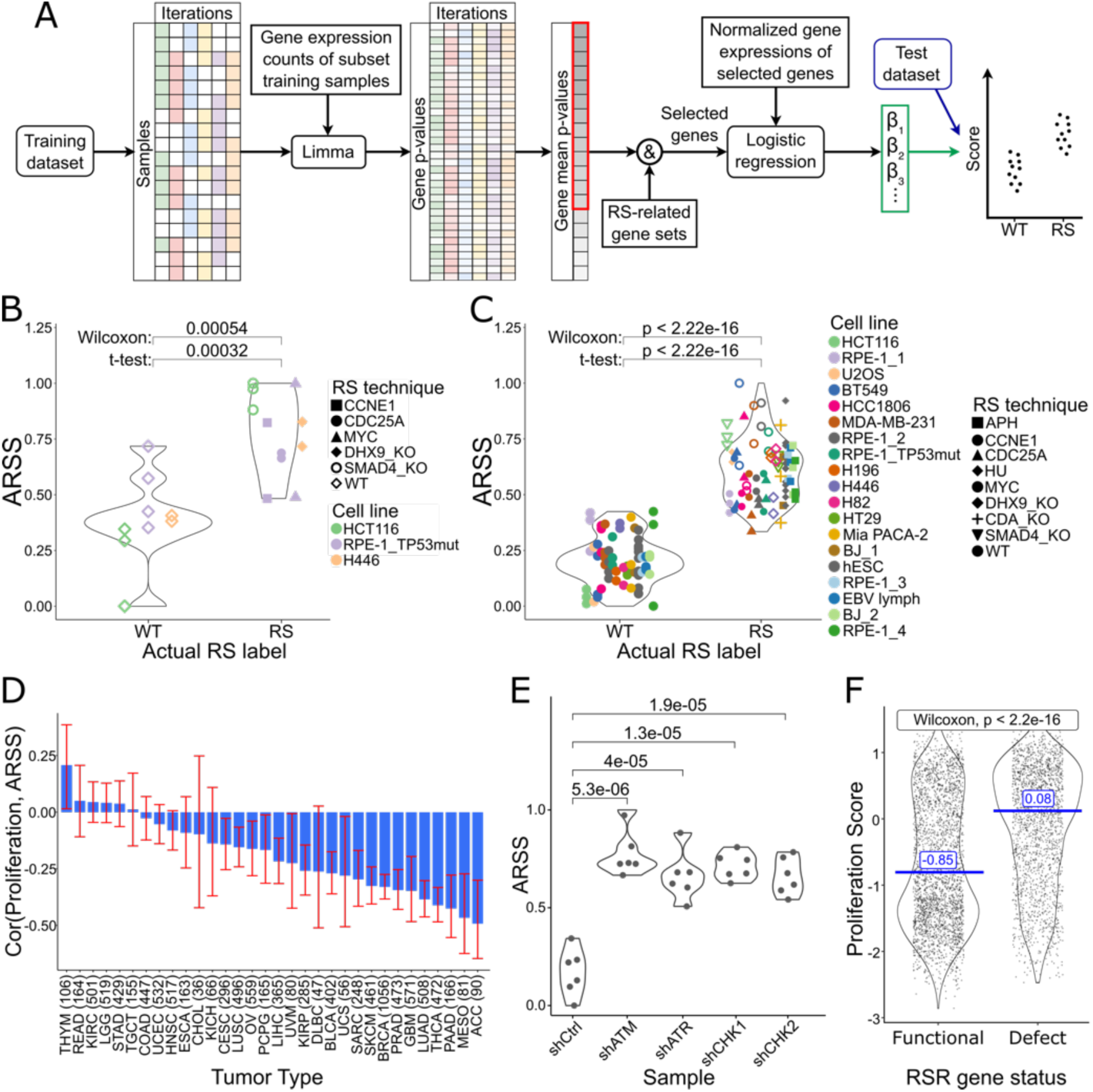
Development and interpretation of a signature of acute replication stress. **(A)** Workflow of the RS prediction model development. Estimating consistently differentially expressed genes using a permutation approach, limma-based differential gene expression determination and calculating gene weights using regularized logistic regression. **(B)** Comparison of the RS scores predicted by a signature for acute RS (ARSS) between samples with (RS) and without (WT) replication stress on an unseen test data set. **(C)** Predictions of the ARSS on the entire dataset combining eleven studies and eight RS induction methods compared to the actual RS status grouped into samples with (RS) and without (WT) replication stress. **(D)** Spearman correlation between the ARSS and a proliferative score (y-axis) in different TCGA tumor types (x-axis). **(E)** Acute RS score in samples with functional RS response (shCtrl) and with silenced gene expression of RS response genes (shATM, shATR, shCHK1, shCHK2). **(F)** Comparison of proliferation between TCGA samples with (Defect) and without (Functional) mutations or copy number losses in RS response genes (ATR, ATM, CHEK1, CHEK2). RS: Replication stress; ARSS: Acute replication stress signature; TCGA: The Cancer Genome Atlas; RSR: Replication stress response

Finally, the model was trained on the entire dataset (Figure 2C; Table 2). Because of the short-term RS-induction of around 48 h, we will call the resulting signature Acute Replication Stress Signature (ARSS).

**Table 2:** Genes in Acute Replication Stress Signature (ARSS) including their weights.

|  | *Gene* | *coef* |
| --- | --- | --- |
| 1 | TFDP2 | 2.596 |
| 2 | SF3A3 | 2.518 |
| 3 | BTG2 | 2.013 |
| 4 | RADX | 1.063 |
| 5 | BRME1 | 1.048 |
| 6 | POLH | 0.930 |
| 7 | SUPT7L | 0.575 |
| 8 | RRM2B | 0.534 |
| 9 | CEP164 | 0.424 |
| 10 | SUN2 | 0.259 |
| 11 | REV3L | 0.254 |
| 12 | AOC2 | 0.193 |
| 13 | PLK1 | -0.039 |
| 14 | PPP2R1A | -0.097 |
| 15 | ERCC1 | -0.097 |
| 16 | VCL | -0.126 |
| 17 | MCM6 | -0.372 |
| 18 | HINFP | -0.421 |
| 19 | H2BC14 | -0.728 |
| 20 | NUCKS1 | -0.861 |
| 21 | POLR1D | -1.527 |
| 22 | ERCC2 | -1.924 |
| 23 | FBXL7 | -2.338 |
| 24 | SPIDR | -2.440 |
| 25 | PIF1 | -2.577 |
| 26 | H2AC14 | -2.885 |
| 27 | AKT3 | -3.735 |
| 28 | PPP2R5A | -4.588 |

Previously, using moderate APH dosages over 100 nM to induce RS has been criticized for halting cell cycle progression through checkpoint activation rather than a physiological response to RS [26]. To assess whether the ARSS, which uses trainings samples with APH levels of 200-400 nM, is physiologically feasible in primary tumors, we compared its association to proliferation. As displayed in Figure 2D, ARSS is negatively correlated with proliferation in TCGA tumor types, contrasting the previous observation of increased proliferation in tumors with RS and genomic instability [20]. To investigate the meaning of the response signature, we used a dataset with defects in the RS response genes *CHK1/2*, *ATR*, *ATM* [24]. Notably, samples with RS response defects (RSRD) displayed significantly higher levels of ARSS (Figure 2E). Defects in checkpoint genes are associated with an increase in proliferation in TCGA primary tumors, as indicated by higher proliferation rates in samples with heterozygous RSRD (Figure 2F), and in cell line experiments with silenced RS-response genes [24,25]. However, disruption of the cell cycle by hydroxyurea exposed an inability to recover from stalled replication forks [24]. This suggests a high level of stress in these RSRD samples that might resemble samples with RS induced by moderate APH dosages as used in ARSS generation.

To explore pathways linking the two signatures we calculated the scores for the ARSS and the RSRD in the TCGA dataset and performed Gene Set Enrichment Analysis [40]. Common enrichment was observed in the *TP53* pathway (q < 0.01 for both signatures) and the gene set for downregulation of the *KRAS* signaling pathway (q < 0.03 for both signatures) [41]. Both describe the inhibition of cell cycle progression at the G1/S checkpoint [42,43], which aligns with the observation of inhibited cell cycle progression in samples treated with moderate APH dosages [26]. These results suggest that the samples with 200-400 nM APH treatment drive the ARSS to reflect conditions of severe RS leading to cells cycle arrest instead of tumorigenic RS.

### A novel RS signature highlights elevated proliferative cell cycle activity in primary tumors versus cell lines

To capture physiological, tumorigenic replication stress, we developed a second signature by excluding samples subjected to severe RS conditions, such as those treated with high concentrations of aphidicolin and hydroxyurea from the dataset (Table 1). This signature was trained using the same methodology (Figure 2A) on the reduced dataset and validated on an unseen test dataset (Figure 3A), confirming the model’s ability to reliably distinguish between RS and non-RS samples. The final model weights (Table 3) were trained on the full dataset and applied to predict RS (Figure 3B).

**Fig 3:**
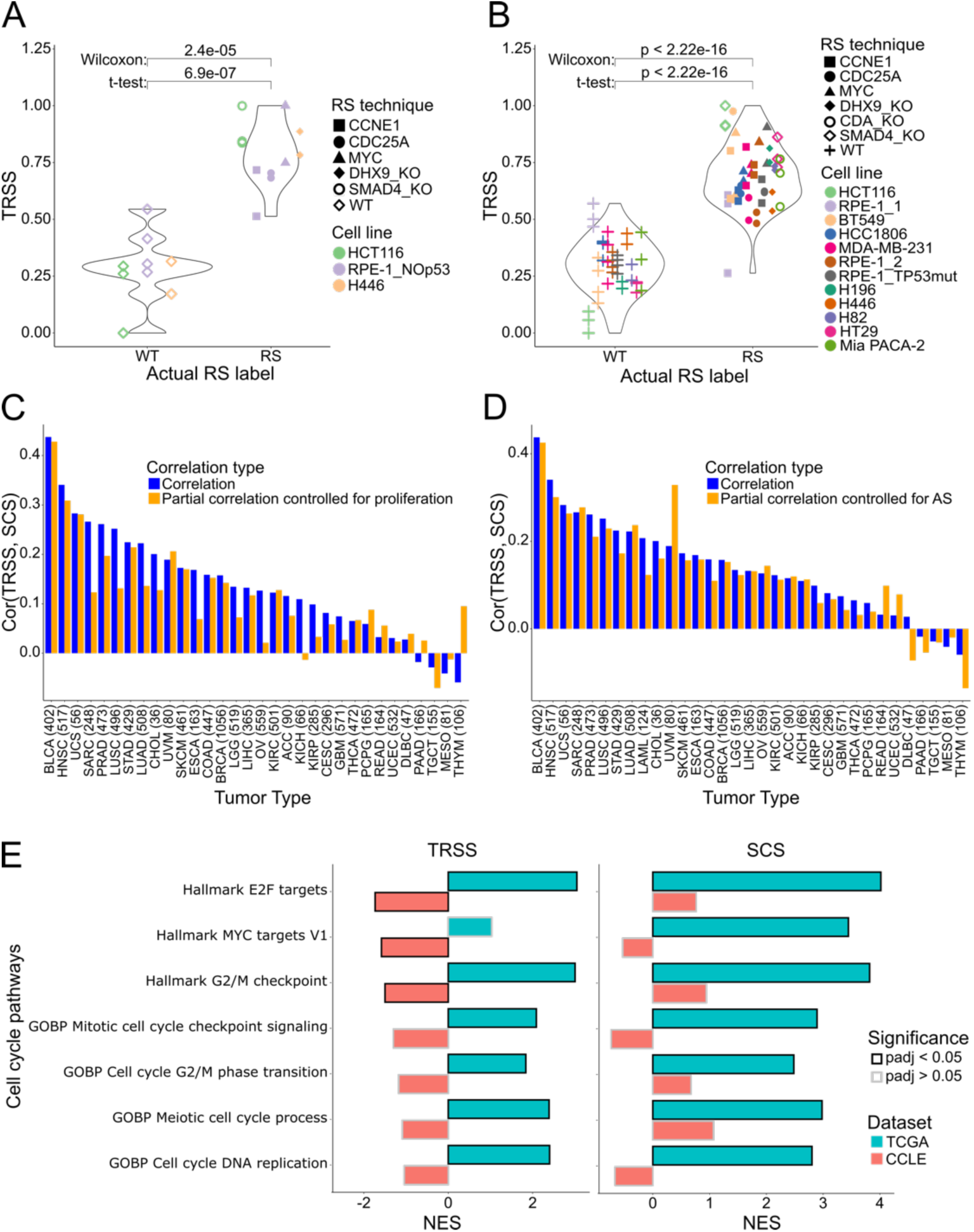
Validation and application of a novel RS signature for tumorigenic RS not confounded by proliferation and aneuploidy. **(A)** Comparison of the RS scores predicted by a signature for tumorigenic RS (TRSS) between samples with (RS) and without (WT) replication stress on an unseen test data set. **(B)** Predictions of the TRSS on the entire dataset combining eleven studies and eight RS induction methods compared to the actual RS status grouped into samples with (RS) and without (WT) replication stress. **(C),(D)** Spearman correlation of the TRSS and the SCS within tumors of TCGA. The y-axis visualizes the non-adjusted (blue) correlation coefficient and the partial (orange) correlation coefficient between repstress score and SCS accounting for proliferation **(C)** and aneuploidy **(D)** in each tumor type (x-axis). **(E)** Gene sets representing regulation of different stages of the cell cycle and their normalized enrichment in association to TRSS (left) and SCS (right) in TCGA (blue) and CCLE (red). RS: Replication Stress; TRSS: Tumorigenic replication stress signature; TCGA: The Cancer Genome Atlas; AS: Aneuploidy score; SCS: Structural Complexity Score; CCLE: Cancer cell line encyclopedia; NES: Normalized enrichment score

**Table 3:** Genes in Tumorigenic Replication Stress Signature (TRSS) including their weights.

|  | *Gene* | *coef* |
| --- | --- | --- |
| 1 | ARID2 | 3.252 |
| 2 | H4C8 | 2.956 |
| 3 | TOP3A | 2.630 |
| 4 | TAF12 | 2.358 |
| 5 | AOC2 | 2.063 |
| 6 | BTG2 | 1.906 |
| 7 | TFDP2 | 1.848 |
| 8 | PHF8 | 1.729 |
| 9 | ATF2 | 1.591 |
| 10 | H2BC12 | 1.415 |
| 11 | H2AC6 | 0.235 |
| 12 | GLI1 | -0.085 |
| 13 | POLD4 | -0.144 |
| 14 | TUBB | -0.390 |
| 15 | BCAM | -0.793 |
| 16 | H4C14 | -1.026 |
| 17 | UFL1 | -1.210 |
| 18 | CMPK2 | -1.329 |
| 19 | FBXL7 | -1.626 |
| 20 | TP53 | -2.693 |
| 21 | PPP2R5A | -3.138 |

To validate the biological relevance of this second RS signature, which we will reference as Tumorigenic RS Signature (TRSS), we examined its association to the SCS in TCGA tumors as well as its dependence on proliferation and aneuploidy using partial correlation. While proliferation had a minor effect on the association (Figure 3C), the confounding by aneuploidy was negligible (Figure 3D). Moreover, we found that the TRSS generally exhibited a weak correlation with proliferation (cor: 0.1) and aneuploidy (cor: 0.07), suggesting the TRSS captures an aspect of genomic instability independent of proliferative capacity or extensive chromosomal changes.

Next, we investigated whether the TRSS reflects the differences in cell cycle activity between primary tumors and cell lines seen in experimental studies. Using GSEA, we calculated the enrichment of cell cycle-related gene sets from the gene ontology biological process [44,45] and hallmark collections [41] dependent on TRSS and SCS. Interestingly, for both scores we found a significant upregulation of various cell cycle gene sets in TCGA primary tumors, whereas this upregulation was not observable in cell lines from the Cancer Cell Line Encyclopedia (CCLE) (Figure 3E; Supplementary Table 4). We asked whether this enrichment implied a cell cycle-related increase of proliferation by examining the deregulation of leading-edge genes associated with the TRSS with a proliferation score and *PCNA* and *MKI67*. Indeed, in TCGA, leading-edge genes of the enriched pathways displayed a higher correlation with the three proliferation measures than non-leading-edge genes as determined by a two-sample Wilcoxon test (all p < 2.2e-16; Supplementary Figure 1A). In cell lines on the other hand, the leading-edge genes indicated a reduction in cell cycle progression (all p < 2.2e-16; Supplementary Figure 1B). The observed increase in cell cycle activity promoting proliferation in primary tumors but not cell lines, captured by the TRSS, hints at an underlying process responsible for the previously discussed paradoxical association of CIN/RS and proliferation.

### Tumorigenic replication stress signature reveals MSH2/6 complex mediated regulation of double-stranded DNA recombination in proteomics data

A recent study in yeast suggests that cellular stress can alter mRNA translation patterns [46]. Using protein expression data from the Clinical Proteomic Tumor Analysis Consortium (CPTAC) database, we investigated proteomic effects of replication stress. Apart from ARID2, which was part of the TRSS signature, two of the proteins most significantly correlated with the TRSS were *MSH6* and *MSH2*, which form the DNA mismatch repair complex MutSα (Figure 4A). *MSH6* protein expression and TRSS exhibited a Spearman correlation of 0.3, while *MSH2* protein expression displayed a correlation of 0.28 (q-value < 2e-14). Interestingly, no other MMR proteins demonstrated a comparably strong association with the TRSS (Supplementary Table 5). The two MutSα proteins were also among the top-ranking proteins in the TCGA proteomics dataset (both q-values < 2e-14), although it only includes about 300 (phospho-)proteins (Supplementary Figure 2A).

**Fig 4:**
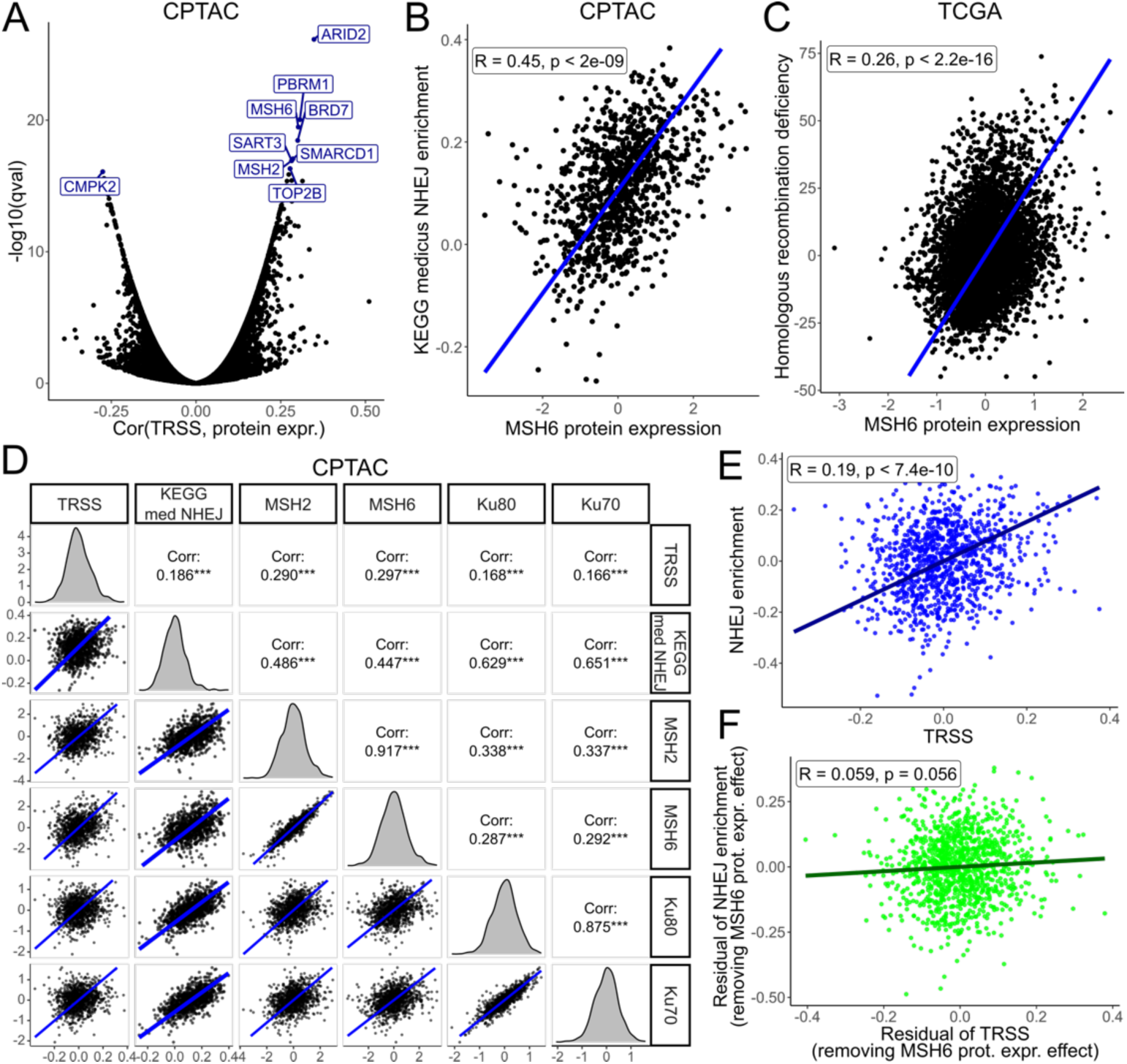
MSH6 protein expression is positively associated with the TRSS and NHEJ, potentially leading to low fidelity DNA repair and genomic scars resembling homologous recombination deficiency. **(A)** Volcano plot visualizing the spearman correlation and q-values of CPTAC protein expressions to the TRSS. **(B)** Activity in the KEGG medicus NHEJ gene set compared to the protein expression of MSH6 in CPTAC. **(C)** Amount of genomic scars typical for homologous recombination deficiencies compared to the protein expression of MSH6 in CPTAC. **(D)** A comparison of TRSS and NHEJ enrichment to protein expression of the proteins in the MSH2/6 complex and the Ku heterodimer in CPTAC. **(E)** Activity in the KEGG medicus NHEJ gene set compared to the TRSS in CPTAC. **(F)** Activity in the KEGG medicus NHEJ gene set compared to the TRSS after adjusting for the effects of MSH6 expression in CPTAC. TRSS: Tumorigenic replication stress signature; CPTAC: Clinical Proteomic Tumor Analysis Consortium; NHEJ: Non-homologous end-joining; TCGA: The Cancer Genome Atlas; KEGG: Kyoto Encyclopedia of Genes and Genomes

Although sensing of mismatches is the primary function of MutSα, its components are also implicated in the DSB repair pathways of NHEJ and homologous recombination [47–49]. Correlation analyses revealed that *MSH2/6* protein expression was significantly associated with single sample GSEA enrichment scores [50] in the KEGG medicus reference NHEJ gene set [51] in CPTAC and TCGA (Figure 4B, Supplementary Figure 2B). *MSH2/6* also correlated with a score quantifying genomic scars resulting from homologous recombination deficiency [52] in TCGA, indicating low fidelity DSB repair (Figure 4C, Supplementary Figure 2C). Further analysis in CPTAC demonstrated that MutSα components as well as the TRSS were associated with Ku70 and Ku80 (Figure 4D), crucial components of the double-strand break repair via NHEJ [53]. The relationship of MutSα and NHEJ also translated into an association between TRSS and NHEJ in CPTAC (Figure 4E, Supplementary Figure 2D). Interestingly, upon partial correlation to control for *MSH2/6* protein expression the association between TRSS and NHEJ activity was removed (Figure 4F, Supplementary Figure 2E), suggesting that the association might be mediated by MutSα.

## Discussion

This study sought to evaluate the proficiencies and limitations of existing RS gene expression signatures and improve upon these limitations to elucidate new insights into the role of replication stress in tumorigenesis.

Using a dataset of labeled samples from the gene expression omnibus, we found that both the repstress score as well as the oncoRS score fail to reliably differentiate RS samples. The repstress score demonstrated an exaggerated dependence on genes associated with cell cycle progression. This is attributable to the selection of *MYC*-associated genes from the G2M-checkpoint and E2F target hallmark pathways, as supported by the strong association to a proliferation score, which also decouples the expected association between the repstress score and a measure for structural aneuploidies. While there are likely situations in which proliferation and replication stress are causally linked [22,23], the general association of RS and genomic instability to proliferation is expected to be weak even in primary tumors, with recent studies suggesting it may further decrease as a result of adaptation to aneuploidies [54,55]. The oncoRS score is only targeted at identifying genes associated with oncogene-induced RS, as acknowledged by the original authors, likely limiting it from generalizing to RS stemming from other sources. The signature may instead over-emphasize other oncogene-specific processes like cell cycle regulation as essential causes of RS [1,12–14]. Methodologically, the score lacks validation of the entire signature, as only one signature gene is explored further. While the present study establishes a link between the oncoRS score and the SCS, both the signature and the SCS rely on copy number alterations in TCGA tumors. The reduced association when controlling for arm-level aneuploidies might indicate that the signature captures aneuploidy-related gene expression, rather than directly measuring RS related to chromosomal instability.

Our first alternative approach leveraged oncogene, gene knockout and drug-based induction of replication stress to detect transcriptional changes and define the ARSS. However, our analysis suggests that the samples with high APH concentrations might contain non-physiological levels of RS, which manifest itself through increased levels of cell cycle arrest as evidenced by the lower proliferation rate in samples with high ARSS. While a slight increase in checkpoint activation might be expected upon DNA damage resulting from RS, the APH samples might represent a gene expression profile of a checkpoint response to excessive DNA replication stress and DNA damage [26]. The good predictive ability on samples with defective RS response indicates that RSRD samples are subjected to large stresses, as is evidenced by McGrail et al.’s observation that RSRD cells are unable to recover from disruption through HU treatment and accumulate in S-phase [24].

The TRSS was therefore designed to minimize these non-physiological biases by excluding samples that might display exaggerated RS. The association of TRSS and SCS across tumor types, independent of proliferation and aneuploidies, suggests that the TRSS reliably captures tumorigenic RS that goes hand in hand with S-CIN. Analysis of point mutations and copy number alterations additionally revealed TRSS connections to genes involved in DNA repair processes, replication fork protection, and the TP53 pathway (Supplemental Material). Furthermore, we found that the difference in proliferative cell cycle activity in primary tumors versus cell lines associated with the TRSS also aligns with the expectations set by the literature, summarized in the introduction, where primary tumors tend to show an increase in proliferation and tumor growth, while cell lines show a slight decrease in cell viability and proliferation upon induction of RS and CIN. This alignment with the literature further indicates that, despite being derived solely from cell line data, the TRSS still generalizes to primary tumors.

Considering this additional evidence for a significant difference in RS-related proliferation between primary tumors and cell lines, a key question arises: Why does RS appear to impair long-term proliferation in cell lines but not in primary tumors? One key difference is the tumor microenvironment, which may be shaped to support the proliferation associated with RS and CIN by (i) reducing the function of T-cells [56], (ii) recruiting pro-tumor M2 macrophages [57,58], and (iii) signaling cancer-associated fibroblasts to provide energy-rich metabolites [59]. Leveraging protein expressions, we found the *MSH2-6* protein complex to be one of the DNA repair components that is most strongly associated with the TRSS. Our findings suggest that this complex might connect RS to an increase in NHEJ activity, potentially exacerbating CIN through low fidelity DNA damage repair. This is consistent with experimental evidence demonstrating that inhibition of *MSH6* leads to a decrease in NHEJ activity [49]. This relationship may be mediated via *MSH6*’s interaction with Ku70 observed in both HeLa cells [49] and oocytes [60]. Another study found that prostate cancers with increased MMR gene expression, including *MSH6*, displayed a higher incidence of deletions at different chromosome loci [61], while NHEJ may add to structural aberrations by generating radial chromosomes or chromothripsis [62]. In line with this Zhang et al. proposed that under RS conditions MutSα may block fork protecting factors causing DNA breaks and chromosomal instability upon misincorporations [63]. Although the extent to which this RS-related NHEJ-activity is linked to CIN requires further investigation, these findings suggest that the TRSS will provide new insights into the role of RS in cancer.

## Methods

### Calculation of oncoRS and repstress scores

The oncoRS [16] and repstress [15] scores were calculated using variance-stabilized transformed gene expressions. OncoRS genes were weighted equally, while repstress genes used the weights defined in the original publication.

### Statistical tests

Correlations were calculated using spearman correlation, and group difference were calculated using Wilcoxon rank-sum tests unless specified otherwise. Where applicable, p-values were adjusted for multiple testing using the Benjamini-Hochberg method.

### Gene Set Enrichment Analysis

Differentially regulated pathways associated with gene signatures were determined using Gene Set Enrichment Analysis using the fgsea package on differentially expressed genes identified via the limmavoom pipeline. Single-sample GSEA was conducted using the gsva package to estimate gene set activity.

### Development of robust gene expression signature

1. RNA sequencing reads were quality-controlled using FASTQC and Trimmomatic and quantified using kallisto.
2. Calculated variance stabilization transformed expression and corrected for batch effects of different RS-inducing treatments and cell lines.
3. Split the dataset into training and test data ensuring independent experiments by stratifying by treatment and cell line.
4. Performed differential gene expression analysis for *n* permutations

a. Sample *p*% from training data.
b. Estimate differential gene expression in RS samples using limma-voom.
c. Save corresponding q-values.
5. Calculate *meanrank* of q-values for differential gene expression.
6. Select genes from RS-adjacent gene sets (Supplementary Table 3).
7. Select the *r* highest ranked genes.
8. Train a regularized logistic regression model with the elastic net coefficient *α*.
9. Validate model on the test set.

The hyperparameters (*n*, *p*, *r*, and *α*) as well as the choice of the mean rank for pre-selection were determined using 10-fold cross validation with the area under the curve (AUC) of the receiver operating characteristic (ROC) curve as performance metric.

## Materials

The accession numbers for gene expressions used for generation and validation of RS signatures are listed in Table 1 & Supplementary Table 2 (accessed: 25. March 2024). Gene expression with RS samples with response defects are available at GSE59227.

Gene sets used were obtained from MsigDB [40,41]: KEGG medicus gene set (MSigdb v2023.2) [51]; Gene Ontology [44,45] and Hallmark (MSigdb v2023.1) [41]

### Omics-data

Gene expression, DNA segment, mutational, and proteomic data in primary tumors generated by The Cancer Genome Atlas (TCGA) Research Network https://www.cancer.gov/tcga [64] were retrieved from the GDC data portal https://portal.gdc.cancer.gov/ (accessed: 03. July 2023).

Gene expression, and proteomic data in primary tumors generated by the National Cancer Institute Clinical Proteomic Tumor Analysis Consortium were retrieved from the PDC portal [65] https://proteomic.data-commons.cancer.gov/pdc/cptac-pancancer (accessed: 10. August 2024).

Gene expression data in cell lines compiled in the Cancer cell line encyclopedia [66] was retrieved through the DepMap data portal https://depmap.org/portal (accessed: 23. August 2023).

### TCGA derivative data

SCS: Distinct and Common Features of Numerical and Structural Chromosomal Instability across Different Cancer Types [18].

Proliferation rate, HRD score, aneuploidy score: The immune landscape of cancer [67] (accessed: 01. July 2024).

Amplification and Deletion calls: Oncogenic Signaling Pathways in The Cancer Genome Atlas [68] (accessed: 27. June 2024).

## Acknowledgments

The authors thank Holger Bastians and Atmika Paul for providing additional RNA sequencing data for SMAD4 knockout-induced replication stress.

## Supporting Captions

Supplementary Information: This file contains a section highlighting genomic alterations associated with the Tumorigenic replication stress signature.

S1 Fig: Association of cell cycle genes to proliferation measures in primary tumors and cell lines.

S2 Fig: Association of the TRSS with MSH2 protein and homologous recombination deficiency.

S3 Fig: Somatic mutations and copy number alterations associated with the TRSS.

S1 Table: Spearman correlation of repstress signature genes to three proliferation measures.

S2 Table: Samples with replication stress-induction by gene amplification or knockout, or treatment with replication stress-inducing drugs for generating a replication stress signature.

S3 Table: Gene sets and genes used as prior knowledge included in the training of the replication stress models.

S4 Table: Tumorigenic Replication Stress Score associated enrichment of cell cycle-related gene sets in the gene ontology biological process (GOBP) and hallmark collections in primary tumors (TCGA) and cell lines (CCLE).

S5 Table: Association of MMR protein expression to the TRSS.

## Notes

### Competing Interest Statement

The authors have declared no competing interest.

